# Spring reproductive success influences autumnal malarial load in a passerine bird

**DOI:** 10.1101/2023.07.28.550923

**Authors:** Romain Pigeault, Camille-Sophie Cozzarolo, Jérôme Wassef, Jérémy Gremion, Marc Bastardot, Olivier Glaizot, Philippe Christe

**Affiliations:** Laboratoire EBI, Equipe EES, UMR CNRS 7267, 86000 Poitiers, France; Department of Ecology and Evolution, Université de Lausanne, Biophore, 1015 Lausanne, Switzerland; Biogéosciences, UMR 6282 CNRS, université de Bourgogne, 6 boulevard Gabriel, 21000 Dijon, France; Muséum cantonal des sciences naturelles - Département de zoologie, Palais de Rumine, Place de la Riponne 6, 1005 Lausanne, Switzerland

**Author notes:** Co-first (Romain Pigeault, Camille-Sophie Cozzarolo).

**Keywords:** avian malaria, annual variations, relapses, recrudescence, recurrences, parasitemia, life history traits, bird

## Abstract

Although avian haemosporidian parasites are widely used as model organisms to study fundamental questions in evolutionary and behavorial ecology of host-parasite interactions, some of their basic characteristics, such as seasonal variations in within-host density, are still mostly unknown. In addition, their interplay with host reproductive success in the wild seems to depend on the interaction of many factors, starting with host and parasite species and the temporal scale under study. Here, we monitored the parasitemia of two haemosporidian parasites – *Plasmodium relictum* (lineage SGS1) and *P. homonucleophilum* (lineage SW2) – in two wild populations of great tits (*Parus major*) in Switzerland over three years, to characterize their dynamics. We also collected data on birds’ reproductive output – laying date, clutch size, fledging success – to determine whether they were associated with parasitemia before (winter), during (spring) and after (autumn) breeding season. Parasitemia of both species dramatically increased in spring, in a way that was correlated to parasitemia in winter. Parasitemia before and during breeding season did not explain reproductive success. However, the birds which fledged the more chicks had higher parasitemia in autumn, which was not associated with their parasitemia in previous spring. Our results tend to indicate that high haemosporidian parasite loads do not impair reproduction in great tits, but high resource allocation into reproduction can leave birds less able to maintain low parasitemia over the following months.

## Introduction

The assumed impact of parasitic infections on animal fitness is at the basis of several evolutionary theories such as the Hamilton-Zuk (Hamilton & Zuk, 1982) or the terminal investment (Stearns, 1989) hypotheses. A recent meta-analysis highlighted the overall negative cost of parasites on reproductive success (Hasik & Siepielski, 2022). This meta-analysis focused on the infection status as a binary variable (parasitized versus non-parasitized), as is the case of many studies. However, the parasite load might better represent the host’s ability to control the infection (resistance) and might be a finer correlate of its physiological or energetical costs (Stjernman et al., 2008; Risely et al., 2018; Sánchez et al., 2018; Methling et al., 2019). A negative association between parasite load and reproductive success was shown in several host-parasite systems (Madsen et al., 2005; Asghar et al., 2011; Gooderham & Schulte-Hostedde, 2011; Hicks et al., 2019; Schoepf et al., 2022) but some studies also show an absence or a positive correlation (Siikamäki et al., 1997; Edler et al., 2004; Raveh et al., 2011; Kulma et al., 2014; Delefortrie et al., 2022).

The different results obtained from studies sometimes involving similar host-parasite pairs could be explained by a number of factors. For instance, the results can vary with the host’s sex. Several studies have reported a negative relationship between parasite load and reproductive success only in males (Sundberg, 1995; Dawson & Bortolotti, 2001; Gooderham & Schulte-Hostedde, 2011; but see Hicks et al., 2019). The age of the infected individuals could also partly blur the signal, since it has been shown that in some species, individuals nearing the end of their lives may invest heavily in reproduction (i.e., terminal investment hypothesis, Velando et al., 2006; Duffield et al., 2018), and in others, that the age-reproductive success relationship follows a bell-shaped curve (Lecomte et al., 2010; Saraux & Chiaradia, 2022). The fluctuation of parasite loads over time may also pose a significant challenge. A large diversity of parasites, ranging from viruses to eukaryotic organisms, exhibit highly dynamic patterns of replication rate, leading to temporal fluctuations of within-host load (e.g., Hasker et al., 2013; Pigeault et al., 2018; Colangeli et al., 2020). These fluctuations can occur on short-term scales, such as daily variations, as well as on long-term scales, spanning months or even years (Martinez-Bakker & Helm, 2015; Prior et al., 2020). Although the parasite load experienced by the host during the breeding period is most likely to have a direct (e.g. pathogenicity, resource exploitation) influence on host’s reproductive success (e.g. Madsen et al., 2005; Asghar et al., 2011; Gooderham & Schulte-Hostedde, 2011; Hicks et al., 2019; Schoepf et al., 2022), the parasite load before the breeding season may also have indirect effects, by influencing for instance premating trade-offs in resource allocation (i.e. carry over effect, Harrison et al., 2011; e.g. Marzal et al., 2013). On the other hand, as life-history theory assumes that components of reproductive effort are costly, investment in reproduction may have longer-term consequences for the host’s ability to clear or at least to control parasite replication rate (Williams, 1966; Stearns, 1989; Sheldon & Verhulst, 1996). This was notably shown in great tits, collared flycatcher and Soay sheeps, in which increased reproductive effort was later associated with higher loads of haemosporidian parasites and strongyle nematodes (Richner et al., 1995, Oppliger et al., 1996, Christe et al. 2012, Nordling et al., 1998; Leivesley et al., 2019). However, the association between natural reproductive effort and parasite load several months after the reproductive event was rarely investigated.

In this study, we used a longitudinal approach to assess whether the parasite load, quantified a few months before and during the breeding season, could explain host reproductive output, and whether reproductive effort could predict parasite load later in the year. To do this, we used avian malaria as a biological system. For over a century, this vector borne disease has been used in studies of host-parasite interactions (Pigeault et al., 2015, Rivero & Gandon, 2018) and provides an excellent model for exploring the influence of parasitic infections on host life-history traits (e.g., Oppliger et al., 1997; Asghar et al., 2015; Pigeault, Cozzarolo, et al., 2018). To date, the vast majority of studies that investigated the influence of malaria infection on bird reproduction used the infection status (i.e., parasitized versus non-parasitized) as a predictor of diverse reproductive parameters (e.g., Sanz et al., 2001; Norte et al., 2009; Podmokła et al., 2014; Zylberberg et al., 2015; Pigeault, Cozzarolo, et al., 2018). The few non-interventional studies that have examined the influence of parasitemia (i.e., quantity of haemosporidian parasites in the peripheral blood of the host) on the reproductive success of birds have reported contrasting results (e.g., Siikamäki et al., 1997; Edler et al., 2004; Asghar et al., 2011; Kulma et al., 2014). On the other hand, drug-induced reduction of parasitemia usually results in higher reproductive success (e.g., Merino et al., 2000; Marzal et al., 2005; Knowles et al., 2010; Schoepf et al., 2022).

A possible explanation for the difference in results between the non-interventional studies and those in which the parasitemia was experimentally reduced is that only its experimental reduction can ensure that treated birds have a permanently lower parasitemia than control individuals. Indeed, avian malaria is characterized by radical temporal variations in parasitemia. After transmission of the parasite by a mosquito, parasitemia increases rapidly and reaches a maximum in 10-20 days (Pigeault et al., 2018; Palinauskas et al., 2018). Activation of the host immune system then controls avian malaria parasite replication rate, but in most cases, it is not able to eliminate it completely, leading to the establishment of the chronic phase of the infection (Valkiūnas, 2005; Asghar et al., 2012) during which parasites persist at low densities for several months or years. However, this chronic phase can be regularly interrupted by recrudescence events when parasitemia increases significantly over a short period of time (three to seven days, Cornet et al., 2014; Pigeault et al., 2023). Consequently, the results of non-interventional studies investigating the relationship between haemosporidian parasitemia and host reproductive success will be highly dependent on the timing of measurements.

Our non-interventional study was carried out over a period of three years, during which field sessions were organized to capture and recapture great tits in order to (i) monitor annual infection dynamics, (ii) study the link between parasitemia and reproductive success, and (iii) investigate the influence of investment in reproduction on parasitemia measured later in the year. Two great tit populations, characterized by different haemosporidian communities – in particular, the most prevalent *Plasmodium* species in each population is not found in the other one – and different overall reproductive parameters (Pigeault, Cozzarolo et al. 2018), were studied here. Although commonly mentioned in the literature, but rarely reported, we predict a significant increase of parasitemia in spring, illustrating spring recurrences (or spring relapses, Applegate, 1971). In light of the studies which showed a trade-off between activation of the great tit’s immune system and their reproduction (e.g., Ots & Hõrak, 1998; Grzędzicka, 2017; Kubacka & Cichoń, 2020), we predict a negative relationship between winter and/or spring parasitemia and reproductive success. Finally, in view of the energy expenditure associated with reproduction (Visser & Lessells, 2001; Nilsson & Råberg, 2001), we also predict that infected individuals who invest substantially in reproduction will later experience higher parasitemia (Hanssen et al., 2005).

## Material and Methods

### Study area and host species

The study was carried out between 2017 and 2019 in two populations of great tits (*Parus major*) breeding in nest-boxes located near the University of Lausanne (Dorigny Forest, 46°31025.60700 N, 6°34040.71400 E, altitude: 380 m) and 15 km apart in the Marais des Monod (46°34019.95300 N, 6°23059.20400 E, altitude: 660 m), both in Switzerland. As in Pigeault, Cozzarolo, et al. (2018), we monitored nest-boxes during each breeding season (from March to June), collecting breeding parameters: laying date, clutch size and fledging success. We trapped breeding great tits in the nest boxes to determine whether they were infected by haemosporidian parasites and, if so, we molecularly identified the parasites involved in the infection (see Appendix 1). We also measured birds’ mass and tarsus length to calculate the scaled mass index of each individual as a proxy for body condition (SMI, Peig & Green, 2009). We categorized the birds’ age as “first-year” or “older” based on whether they already had their first post-nuptial molt or not, by looking at their wing feathers. To correct for between-year differences in average breeding time, laying date was standardized by using the first year of the study as a reference (2017, 1 April = day 1). The differences in laying date between 2017 and each subsequent year were subtracted from the actual laying date of that particular year (see Allander & Bennett, 1995).

In addition to the spring monitoring, birds were also captured with mist nets during winter (between January and end of March) and autumn (between mid-October and mid-December). Three capture sessions were done in winter and in autumn in the Dorigny Forest, while for logistic reasons it was only possible to perform two sessions per season in the Marais des Monod. We also took blood samples and morphometric measurements on all caught individuals. No capture session was carried out in summer because we know from previous attempts that it is very difficult to capture great tits during this period in our study area.

### Molecular analyses

Only birds captured at least twice in a focal year and diagnosed as infected by *Plasmodium* parasite on at least one of their capture dates were retained for analyses (see **Appendices, section 1** for diagnosis and parasite identification protocol). The quantification of parasitemia in all the blood samples was carried out using qPCR (see **Appendices, section 2**). Parasitemia was calculated with relative quantification values (RQ). RQ can be interpreted as the fold‐amount of target gene (*Plasmodium* 18S rDNA) with respect to the amount of the reference gene (avian 18S rDNA) and is calculated as 2^‐(Ct_*Plasmodium* – Ct_bird)^. For convenience, RQ values were standardized by ×10^4^ factor. Since some individuals were trapped and blood sampled several times per season, we calculated the average parasitemia of individuals for each season when we wanted to compare it between seasons.

### Statistical analysis

We used a mixed model procedure with a normal error structure and bird individual fitted as a random factor to test for an effect of month of capture on change in parasitemia (‘lme4’ package, Bates et al., 2014). Sex, age, *Plasmodium* lineage and year were added as fixed factors into the model. As parasitemia could be expected to be a non-linear function of month of capture, because of spring recurrences, the quadratic term month of capture was added to assess whether it significantly improved the model fit. In order to study the influence of the parasitemia measured in winter and spring on the reproductive parameters and the impact of reproductive parameters on the parasitemia measured in autumn, we calculated the average parasitemia of individual birds within each season of each year. In fact, some individuals were captured several times in the winter or in autumn of a focal year, while others were captured only once, so we used average seasonal parasitemia to reduce data complexity. Then, for a focal year, we used generalized linear models (glm), with an error structure appropriate to each response variable (see **Appendices, Table S1**), to study the impact of winter and spring parasitemia on bird’s reproductive parameters. We also used a glm to evaluate the influence of the birds’ reproductive parameters on their parasitemia measured in the autumn. Previous studies showed that co-infection by at least two different haemosporidian parasite genera was highly prevalent in the two great-tits populations monitored here (Pigeault, Cozzarolo, et al., 2018). As co-infection with *Plasmodium* and *Leucocytozoon* may affect life history traits of great tits (see Figure 2 in Pigeault, Cozzarolo, et al., 2018), birds were also screened for *Leucocytozoon* infection (see **Appendix 1, section 1**). However, detection of *Leucocytozoon* was not achieved for 12 individuals. Given the relatively small size of our dataset and the fact that including *Leucocytozoon* infection status as an explanatory variable did not change the conclusions of our study, all the analyses with *Leucocytozoon* infection status fitted as an explanatory variable are presented in the Appendix (Section 3).

All statistical analyses were carried out using the R statistical software (v. 4.1.3). The raw data and the R script used for the analyses and to produce the figures are available on the figshare repository (10.6084/m9.figshare.23695422).

### Ethics statement

This study was approved by the Ethical Committee of the Vaud Canton veterinary authority (authorization number is 1730.4).

## Results

### Variations in parasitemia

We followed the annual infection dynamics in 70 individuals (31 females and 39 males). We detected four lineages of *Plasmodium* [BT7 (*Plasmodium* sp.): 2, TURDUS1 (*P. circumflexum*): 4, SW2 (*P. homonucleophilum*): 20, SGS1 (P. *relictum*): 43] and one lineage of *Haemoproteus* [PARUS1 (*H. majoris*): 1] but we focused our subsequent analyses on the two most prevalent lineages (i.e. SW2 and SGS1). The parasite communities were very different in both sites. The SW2 lineage was only observed in the Marais des Monod while hosts infected by SGS1 were only captured in the Dorigny forest.

A significant influence of month of capture on the parasitemia was observed but only when month was added as a quadratic term in the model (model 1, month: χ2 = 62.17, p < 0.0001, month^2^: χ2 = 63.205, p < 0.0001, Figure 1). We indeed observed that the parasitemia followed a bell-shaped function: peaking during spring and decreasing thereafter (**Figure 1**). When spring captures were removed from our analyses, we no longer observed an influence of month on parasitemia (model 2, month: χ2 = 1.21, p = 0.271, month^2^: χ2 = 1.09, p = 0.315).

**Figure 1.**
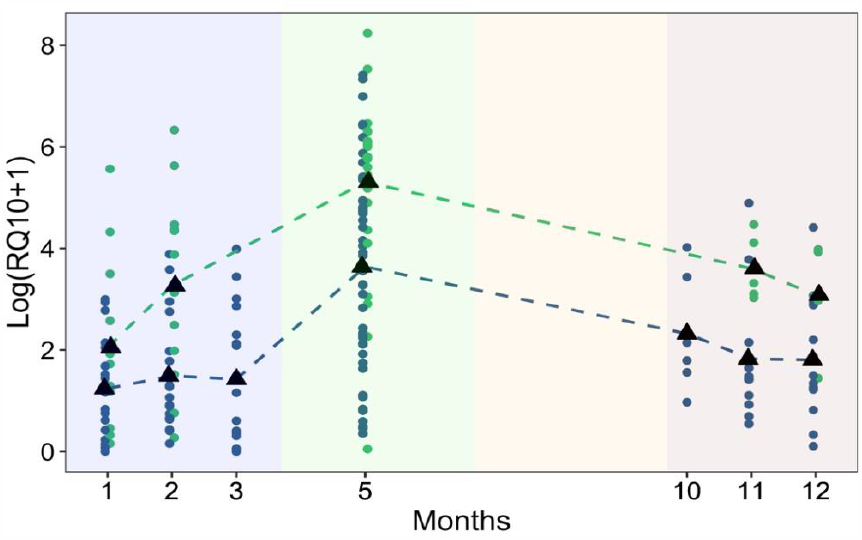
Annual variation of avian malaria within-host infection load. Each point represents the parasitemia (Log RQ10+1) of an individual caught at least twice during a same year. Blue dots correspond to individuals infected with *P. relictum* SGS1, green dots correspond to individuals infected with *P. homonucleophilum* SW2. The colored rectangles represent the seasons. Blue: winter, green: spring, yellow: summer, light red: autumn. Months: 1 = January, 2 = February, 3 = March, 5 = May, 10 = October, 11 = November, 12 = December.

**Figure 2.**
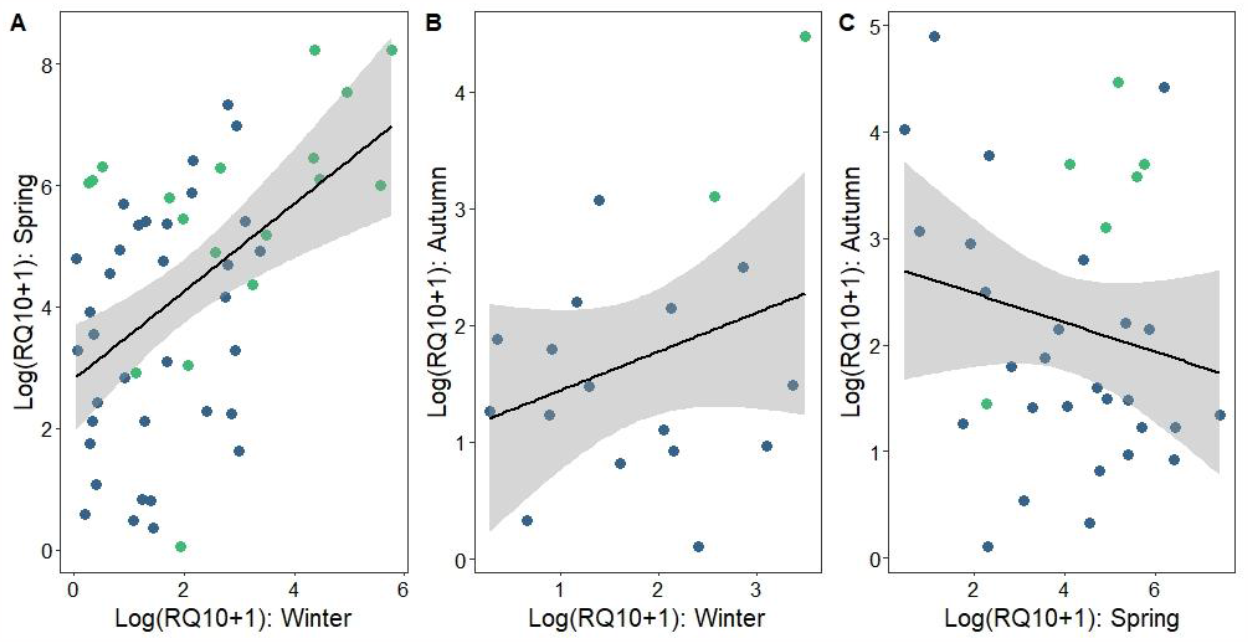
Relationship between parasitemia measured at different season. (A) Relationship between parasitemia in spring and winter, (B) autumn and winter and (C) autumn and spring. Blue dots correspond to individuals infected with *P. relictum* SGS1, green dots correspond to individuals infected with *P. homonucleophilum* SW2.

Although the general shape of infection for both lineages was similar, we observed that host infected by SW2 showed higher parasitemia than those infected by SGS1 (model 1, χ2 = 31.69, p < 0.0001, mean RQ10 ± se, SW2 = 294 ± 99, SGS1 = 148 ±87, Figure 1). Parasitemia differed between years (model 1, χ2 = 9.44, p = 0.002, mean RQ10 ± se, 2017 = 174 ± 69, 2018 = 83 ± 31, 2019 = 247 ± 145) but we did not observe any difference between host sex and age (model 1, sex: χ2 = 1.41, p = 0.235, age: χ2 = 0.002, p = 0.967).

After calculating the average parasitemia of hosts within each season, we observed a significant relationship between winter and spring parasitemia (model 3, χ2 = 6.16, p = 0.010). The most infected hosts in winter exhibited higher parasitemia in spring (**Figure 2**). Conversely, we did not observe any relationship between winter and autumn parasitemia or between spring and autumn (model 4-5, χ2 = 0.62, p = 0.445, χ2 = 0.04, p = 0.837, respectively, **Figure 2**). We could not look for a correlation between parasitemia in autumn and subsequent winter, because of the low sample size (N = 2). As observed previously in the analysis of the annual dynamics of *Plasmodium* infection, within each season, the parasitemia was higher in individuals infected by SW2 than in birds infected by SGS1 (model 3: χ2 = 8.40, p = 0.006, model 4: χ2 = 4.77, p = 0.049, model 5: χ2 = 6.55, p = 0.016).

### Relationship between parasitemia and reproductive parameters

Parasitemia recorded during the winter period did not influence the laying date (model 6, χ2 = 0.244, p = 0.623), the investment in the reproduction or the reproductive success of the hosts (model 7, clutch size: χ2 = 0.103, p = 0.748, model 8, number of fledged chicks: χ2 = 0.479, p = 0.489). We also did not observe any relationship between parasitemia measured during the reproductive period and any reproductive parameters (model 9-11, laying date: χ2 = 0.037, p = 0.847, clutch size: χ2 = 0.141, p = 0.707, number of fledged chicks: χ2 = 0.001, p = 0.975). However, the number of fledged chicks was higher in hosts infected by SW2 than in birds infected by SGS1 (model 11, χ2 = 7.60, p = 0.006, mean ± se, SW2 = 6.22 ± 0.46, SGS1 = 4.20 ± 0.32). Finally, the clutch size did not impact the parasitemia measured later in autumn (model 12, χ2 = 1.340, p = 0.247, **Figure 3A**), but we observed a significant effect of the number of fledged chicks (model 12, χ2 = 5.91, p = 0.015). Individuals with high fledging success also had higher parasitemia the following autumn (**Figure 3B**). We observed a negative relationship between individual SMI and autumn parasitemia (model 12, χ2 = 4.66, p = 0.031). The birds with the lowest body condition were those that had the highest parasitemia (**Figure S1**). However, there was no relationship between the breeding success (i.e., the number of fledged chicks) of birds in spring and their body condition a few months later in autumn (model 13, χ2 = 0.55, p = 0.457).

**Figure 3.**
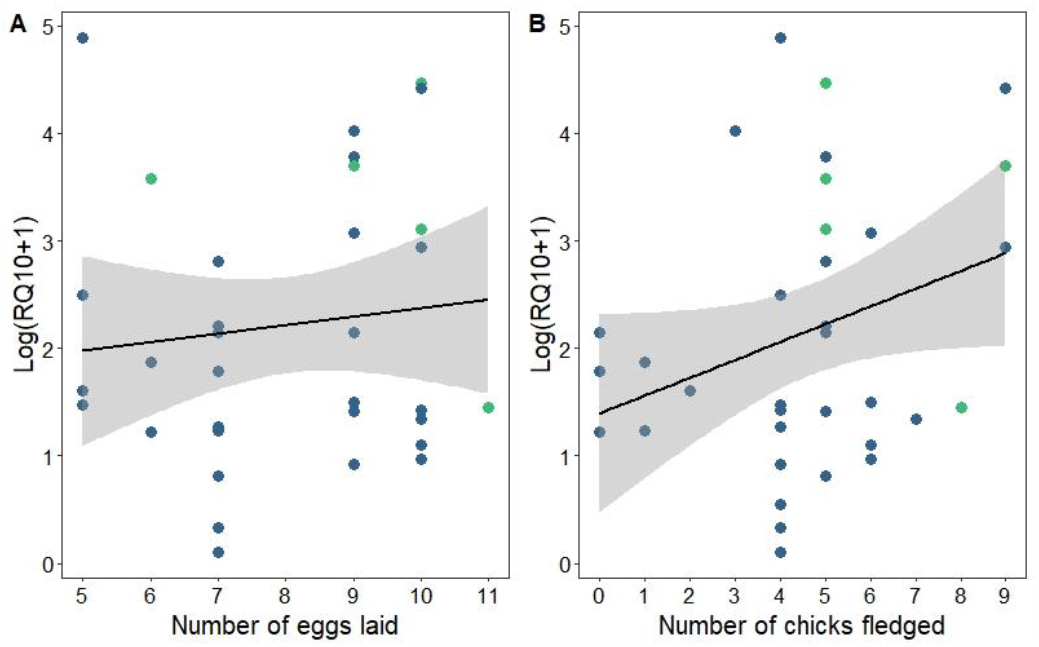
Influence of reproductive parameters monitored in spring and parasitemia measured a few months later in autumn. Relationship between **(A)** the number of eggs laid or **(B)** the number of chicks fledged and parasitemia (Log RQ10+1). Blue dots correspond to individuals infected with *P. relictum* SGS1, green dots correspond to individuals infected with *P. homonucleophilum* SW2.

## Discussion

Using data collected from two wild populations of great tits infected by two different *Plasmodium* lineages, we highlighted that parasitemia increased drastically between winter and spring, and we found that increased reproductive allocation in spring is associated with higher parasitemia in autumn.

Numerous studies have attempted to characterize the yearly fluctuations of infection prevalence in different bird populations. They either found no seasonality in probability of infection (Himalayan bird community; Ishtiaq et al., 2017), a link with migration (Hellgren et al., 2013; Pulgarín-R et al., 2019), or seasonality patterns that vary across *Plasmodium* lineages (Neto et al., 2020) and geographical regions (Cosgrove et al., 2008; Lynton-Jenkins et al., 2020; Neto et al., 2020). On the other hand, little is known about annual within-host variation in parasitemia. In a wild bird community in Slovakian woodlands, *Plasmodium* sp. parasitemia peaked in summer (Šujanová et al., 2021) while *Plasmodium relictum* parasitemia in captive house sparrows (*Passer domesticus*) in Spain varied monthly without clear seasonal pattern (Garcia-Longoria et al., 2022). In the present study, we report that, for two *Plasmodium* lineages, the parasitemia followed a bell-shaped function. It increased more than tenfold between winter and spring, and then decreased in autumn to winter levels. Our study illustrates a phenomenon described by Applegate & Beaudoin in the 70s but rarely documented since: spring recurrences (or spring relapses Applegate, 1970). Spring recurrences can be due to several factors likely acting on bird’s physiology or immunity, such as resource availability (Cornet, Bichet, et al., 2014), environmental stressors (Becker et al., 2020; Pigeault et al., 2023), immune challenges, co-infections with other parasites (Palinauskas et al., 2011; Reinoso-Pérez et al., 2020; Garcia-Longoria et al., 2022) and energy allocation to other functions such as reproduction (Christe et al., 2012).

Although various factors may be responsible for triggering recurrences of infection, we have demonstrated that the spring parasitemia of birds was positively correlated to their parasitemia recorded in winter. However, contrary to our predictions, we did not observe any effect of both winter and spring parasitemia on bird reproductive parameters. Indeed, individuals with extremely high parasitemia (RQ > 1000) did not lay more eggs nor fledged more chicks than those with very low parasitemia (RQ < 1). This result is not consistent with a premating trade-off in resource allocation (i.e., carry-over effect) or with a reallocation of host resources towards immunity during the mating period (Stearns, 1989; Harrison et al., 2011; Stahlschmidt et al., 2013; Albery et al., 2020).

Interestingly, while the parasitemia in winter is a significant predictor of the intensity of the spring recurrence, we did not observe any relationship between parasitemia measured in winter or in spring and parasitemia recorded the following autumn. This result suggests that between spring and autumn, biotic and/or abiotic parameters may have modified the interactions between *Plasmodium* and its host. Infections by new parasites during spring or summer periods could directly or indirectly influence the within-host infection dynamics of *Plasmodium* (Cellier-Holzem et al., 2010). Our study was not designed to test this hypothesis, but we noted that the birds recaptured in the autumn were all infected with the same lineage of *Plasmodium* that was identified in the spring. Further, we did not record any new haemosporidian parasite infection. However, we cannot exclude the possibility that, during the breeding season, the birds were infected by other parasites (e.g. gastrointestinal nematodes, viruses; regarding co-infections with other *Leucocytozoon*, see **Appendices, section 3**).

Inter-individual variability in reproductive investment may also explain why there was no relationship between the parasitemia measured at the beginning of the reproductive period and that measured several months later in autumn. Because immunity and reproduction compete for host resources, in resource-limited environments, hosts that reproduce should have fewer resources to allocate to immune defense which may ultimately influence within-host infection dynamics. Increased allocation to reproduction was found to be associated with increased load of gastrointestinal nematodes in wild Soay sheep during both late gestation and early lactation (Leivesley et al., 2019). Brood manipulation studies on birds showed that increased allocation to reproduction was associated with greater parasite loads and less effective immune responses at the end of the breeding period (Richner et al., 1995, Oppliger et al., 1997, Knowles et al., 2009; Christe et al., 2012). Here, we observed birds with the highest reproductive success tended to be those with the highest parasitemia 6-8 months later. Although supported by a small sample size (n = 36), this result suggests a long-term effect of investment in reproduction on the ability of hosts to control the replication rate of blood parasites.

Finally, in addition to seasonal variations, we observed a significant difference in the parasitemia in birds, depending on the *Plasmodium* lineage involved in the infection. Hosts infected with *P. homonucleophilum* SW2 had a higher parasitemia than hosts infected with *P. relictum* SGS1, irrespective of the time of year when the blood samples were taken. This result is consistent with that of an earlier study conducted on the same great tits populations in 2009-2011 (Rooyen et al., 2013). Interestingly, we also found that birds infected by *P. homonucleophilum* showed higher reproductive success than those infected by *P. relictum*. The fact that *P. relictum* SGS1 is a very generalist and widespread lineage (147 host species and worldwide distribution according to MalAvi database; Bensch et al. 2009) compared to the relatively less common *P. homonucleophilum* SW2 (33 host species, found only in Afro-Eurasia) might correlate with these differences, in line with the hypothesis of a trade-off between generalism and virulence in parasites (Leggett et al. 2013). Peak parasitemia (a proxy for virulence) was negatively correlated with host breadth (a proxy for generalism) in primate malaria parasites (Garamszegi 2006), but, to our knowledge, there is no such evidence for avian haemosporidians. However, there does not seem to be a trade-off between generalism and prevalence in avian *Plasmodium* (Hellgren et al. 2009). Alternatively, *P. relictum* might also have been infecting the Dorigny population for longer than *P. homonucleophilum* has been infecting the Monod population; if this is the case, birds may have had more time to evolve specific immune defenses to limit the virulence of *P. relictum*. Our data do not allow to determine whether the differences observed between hosts infected by these two *Plasmodium* species are the result of differences inherent to the two blood parasites, of the vertebrate host genetic background, of their environment, or interactions between all these factors. Indeed, the individuals infected with *P. homonucleophilum* all originated from the Monod marshlands, whereas the birds infected with *P. relictum* all came from the Dorigny forest. Although these two populations are only 15km apart, we have never observed birds migrating between the two populations and the habitats are very different (large marshy forest massif *versus* periurban forest patches on the campus of the University of Lausanne, respectively).

In conclusion, our monitoring of haemosporidian parasitemia in great tits across seasons evidenced a spring recurrence, the triggers of which are still to be determined. Further, our study supported the idea that strong allocation in reproduction incurs costs later in life, without evidence that higher parasitemia prior to and during breeding season reduce reproductive success. Interestingly, we found this pattern in two great tit populations, infected by two different *Plasmodium* species.

## Supplementary Information

Appendices include:

Section 1: Haemosporidian parasite detection

Section 2: Quantification of parasitemia

Section 3: Results of the analyses with *Leucocytozoon* infection status fitted as explanatory variable

Table S1: Description of the statistical models presented in the main text.

Figure S1: Relationship between scale mass index and parasitemia in birds captured in autumn

## Data, scripts, code, and supplementary information availability

Data and scripts are available online: https://doi.org/10.6084/m9.figshare.23695422. Supplementary information can be found on the bioRxiv preprint server in the Supplementary Material section under this link: https://doi.org/10.1101/2023.07.28.550923.

