## Appendix for "Spring reproductive success influences autumnal malarial load in a passerine bird"

* Co-first

**Section 1: Haemosporidian parasite detection**

**Section 2: Quantification of infection intensity**

**Section 3: Results of the analyses with *Leucocytozoon* infection status fitted as explanatory variable**

**Table S1: Description of statistical models presented in the main text.**

**Figure S1: Relationship between the scale mass index and the intensity of infection of birds captured in autumn.**

### Section 1: Haemosporidian parasite detection

Haemosporidian parasite detection was performed as in Pigeault, Cozzarolo et al. (2018). Briefly, blood samples were collected from the medial metatarsal vein (2-3uL). A nested PCR targeting a fragment of cytochrome b gene of mitochondrial genome of the haemosporidian parasites (i.e. *Haemoproteus, Leucocytozoon* and *Plasmodium,* Hellgren et al., 2004) was performed in triplicates on all samples after DNA was extracted from blood using a DNeasy Blood & Tissue Kit (Qiagen, Switzerland) following the manufacturer's instructions. Amplification products were visualized on agarose gels after electrophoresis to select infected samples. Positive samples were sequenced via Sanger sequencing and haemosporidian lineages were identified by searching the sequences in the MalAvi database (http://130.235.244.92/Malavi/, Bensch et al., 2009). We considered double peaks on DNA chromatographs as an indication for the presence of more than one lineage in a sample. As *Plasmodium* and *Haemoproteus* gene fragments were amplified with the same primer pair, we were not able to distinguish between multiple *Plasmodium*, multiple *Haemoproteus* or *Plasmodium* + *Haemoproteus* infections. For this reason, we were not able to include co-infection with *Haemoproteus* as a covariate in our analyses.

Bensch, S., Hellgren, O., & Pérez-Tris, J. (2009). MalAvi: a public database of malaria parasites and related haemosporidians in avian hosts based on mitochondrial cytochrome b lineages. *Molecular Ecology Resources*, **9**(5), 1353-1358. https://doi.org/10.1111/j.1755-0998.2009.02692.x

Pigeault R, Cozzarolo C-S, Choquet R, Strehler M, Jenkins T, Delhaye J, Bovet L, Wassef J, Glaizot ,Christe P (2018) Haemosporidian infection and co-infection affect host survival and reproduction in wild populations of great tits. *International Journal for Parasitology*, 48, 1079–1087. https://doi.org/10.1016/j.ijpara.2018.06.007

Hellgren, O, Waldenström, J, Bensch, S.(2004) A new PCR assay for simultaneous studies of Leucocytozoon, Plasmodium, and Haemoproteus from avian blood. *Journal of Parasitology*, **90**(4), 797-802. https://doi.org/10.1645/GE-184R1

### Section 2: Quantification of parasitemia

The quantification of parasitemia was carried out using qPCR with a protocol adapted from Christe et al., 2011. DNA was extracted from the blood using a standard protocol (Qiagen DNeasy 96 blood and tissue kit). For each individual, we conducted a multiplex qPCR. We targeted the nuclear *cyt b* gene of *Plasmodium* (Primers: L4050Plasmo: 5′-GCTTTATGTATTGTATTTATAC-3′, H4121Plasmo: 5′-GACTTAAAAGATTTGGATAG-3′, TaqMan probe: CY3-CYTb-BHQ2: 5′- CCTTTAGGGTATGATACAGC-3′) and the 18s rRNA gene of the bird (Primers: Plasmo18S-f: 5′-GGCAGCTTTGGTGACTCTAGA -3′, Plasmo18S-r: 5′-AGTTGATAGGGCAGACATTCG-3′, TaqMan probe: FAM-18S-BHQ1: 5′-AACCTCGAGCCGATCGCACG -3′). All samples were run in triplicate (Bio-Rad CFX96TM Real-Time System). Parasite number was calculated with relative quantification values (RQ). RQ can be interpreted as the fold-amount of target gene (*Plasmodium* 18s rDNA) with respect to the amount of the reference gene (Bird18s rDNA) and are calculated as $2^{-(Ct18s\ \mathit{Plasmodium}\ -\ Ct18s\ Bird)}$. For convenience, RQ values were standardized by $\times 10^4$ factor and log-transformed (log(x, base = exp(1))).

Christe P, Glaizot O, Strepparava N, Devevey G, Fumagalli L (2011) Twofold cost of reproduction: an increase in parental effort leads to higher malarial parasitemia and to a decrease in resistance to oxidative stress. *Proceedings of the Royal Society B: Biological Sciences*, **279**(1731), 1142-1149. https://doi.org/10.1098/rspb.2011.1

### Section 3: Results of the analyses with *Leucocytozoon* infection status fitted as explanatory variable

**Table 1: response variable : bird parasitemia (whatever the month of capture)**

| Explanatory variables | AIC | LRT | p-value |
|---|---|---|---|
| year | 677.56 | 9.397 | 0.002174 |
| *Plasmodium* lineage | 697.22 | 29.050 | 7.052e-08 |
| sex | 670.84 | 2.676 | 0.101877 |
| age | 668.38 | 0.211 | 0.646262 |
| month | 729.55 | 61.385 | 4.693e-15 |
| *Leucocytozoon* infection status (0/1) | 668.66 | 0.499 | 0.480141 |
| I(month^2) | 731.34 | 63.174 | 1.892e-15 |

**Table 2: response variable : spring parasitemia**

| Explanatory variables | AIC | F | p-value |
|---|---|---|---|
| year | 209.23 | 0.9791 | 0.327952 |
| *Plasmodium* lineage | 217.12 | 8.5010 | 0.005618 |
| sex | 212.36 | 3.8164 | 0.057278 |
| age | 208.10 | 0.0000 | 0.998504 |
| RQ10_Winter | 212.92 | 4.3505 | 0.042968 |
| *Leucocytozoon* infection status (0/1) | 209.10 | 0.8621 | 0.358322 |

**Table 3: response variable : fall parasitemia**

| Explanatory variables | AIC | F | p-value |
|---|---|---|---|
| year | 51.127 | 0.0737 | 0.79158 |
| *Plasmodium* lineage | 56.556 | 3.8641 | 0.07769 |
| sex | 52.933 | 1.2031 | 0.29841 |
| age | 51.002 | 0.0000 | 0.99966 |
| RQ10_Winter | 51.741 | 0.4442 | 0.52019 |
| *Leucocytozoon* infection status (0/1) | 51.018 | 0.0096 | 0.92401 |

**Table 4: response variable : laying date**

| Explanatory variables | AIC | Chisq | p-value |
|---|---|---|---|
| RQ10_Spring | 415.30 | 0.3046 | 0.58102 |
| *Plasmodium* lineage | 415.83 | 0.8398 | 0.35944 |
| SMI | 416.30 | 1.3036 | 0.25355 |
| age | 419.79 | 4.7921 | 0.02859 |
| sex | 415.00 | 0.0031 | 0.95577 |
| year | 420.00 | 5.0100 | 0.02520 |
| *Leucocytozoon* infection status (0/1) | 417.69 | 2.6931 | 0.10078 |

**Table 5: response variable : clutch size**

| Explanatory variables | AIC | Chisq | p-value |
|---|---|---|---|
| RQ10_Spring | 273.85 | 0.14638 | 0.7020 |
| *Plasmodium* lineage | 274.84 | 1.13872 | 0.2859 |
| SMI | 273.93 | 0.23307 | 0.6293 |
| age | 274.03 | 0.32598 | 0.5680 |
| sex | 273.93 | 0.22514 | 0.6352 |
| year | 273.89 | 0.18611 | 0.6662 |
| *Leucocytozoon* infection status (0/1) | 273.74 | 0.03749 | 0.8465 |

**Table 6: response variable : number of chicks fledged**

| Explanatory variables | AIC | Chisq | p-value |
|---|---|---|---|
| RQ10_Spring | 307.65 | 0.2300 | 0.63153 |
| *Plasmodium* lineage | 315.89 | 8.4700 | 0.00361 |
| SMI | 307.75 | 0.3211 | 0.57092 |
| age | 309.27 | 1.8488 | 0.17392 |
| sex | 307.49 | 0.0655 | 0.79798 |
| year | 309.10 | 1.6773 | 0.19529 |
| *Leucocytozoon* infection status (0/1) | 308.58 | 1.1519 | 0.28316 |

**Table 7: response variable = fall parasitemia**

| Explanatory variables | AIC | F | p-value |
|---|---|---|---|
| year | 99.514 | 4.4114 | 0.04739 |
| sex | 94.623 | 0.5570 | 0.46335 |
| age | 94.022 | 0.1236 | 0.72846 |
| clutch_size | 94.116 | 0.1909 | 0.66642 |
| fled_success (number of chicks fledged) | 97.380 | 2.6547 | 0.11748 |
| *Plasmodium* lineage | 98.714 | 3.7388 | 0.06613 |
| SMI | 98.539 | 3.5940 | 0.07120 |
| *Leucocytozoon* infection status (0/1) | 93.921 | 0.0514 | 0.82271 |

**Table S1: Description of statistical models presented in the main text.** "N" gives the number of birds included in each analysis. "Maximal Model" includes the complete set of explanatory variables. "Minimal model" gives the model containing only the significant variables and their interactions. Brackets indicate variables fitted as random factors. Curly brackets indicate the error structure used (n: normal errors, p: Poisson). Plsm: *Plasmodium*, SMI: scaled mass index.

| Variable of interest | Resp. variable | Model nb | N | Maximal model | Minimal model | R subroutine {error struct} |
|---|---|---|---|---|---|---|
| Parasite load | Log(RQ10+1) | 1 | 192 | year_of_capture + Plsm_lineage * sex * age * month_of_capture + I(Month_Numbers^2) + (1\|Ring) | year_of_capture + Plsm_lineage + month_of_capture + I(Month_Numbers^2) + (1\|Ring) | lmer {n} |
| Parasite load | Log(RQ10+1) | 2 | 121 | year_of_capture + Plsm_lineage * sex * age * month_of_capture + I(Month_Numbers^2) + (1\|Ring) | year_of_capture + Plsm_lineage + (1\|Ring) | lmer {n} |
| Average parasite load in spring | Log(RQ10+1) | 3 | 53 | year_of_capture + Plsm_lineage * sex * age * RQ_Winter | year_of_capture + Plsm_lineage + RQ_Winter | glm {n} |
| Average parasite load in autumn | Log(RQ10+1) | 4 | 18 | year_of_capture + Plsm_lineage * sex * age * RQ_Winter | Plsm_lineage | glm {n} |
| Average parasite load in autumn | Log(RQ10+1) | 5 | 36 | year_of_capture + Plsm_lineage * sex * age * RQ_Spring | year_of_capture + Plsm_lineage | glm {n} |
| Laying date | day | 6 | 53 | year_of_capture + Plsm_lineage * age * sex * SMI_Winter* RQ_Winter | 1 | glm {n} |
| Number of eggs laid | eggs | 7 | 53 | year_of_capture + Plsm_lineage * age * sex * SMI_Winter* RQ_Winter | 1 | glm {p} |
| Number of chicks fledged | Chicks | 8 | 53 | year_of_capture + Plsm_lineage * age * sex * SMI_Winter* RQ_Winter | 1 | glm {p} |
| Laying date | day | 9 | 71 | year_of_capture + Plsm_lineage * age * sex * SMI_Winter* RQ_Winter | 1 | glm {n} |
| Number of eggs laid | eggs | 10 | 71 | year_of_capture + Plsm_lineage * age * sex * SMI_Winter* RQ_Winter | 1 | glm {p} |
| Number of chicks fledged | Chicks | 11 | 71 | year_of_capture + Plsm_lineage * age * sex * SMI_Winter* RQ_Winter | Plsm_lineage | glm {p} |
| Average parasite load in autumn | Log(RQ+1) | 12 | 36 | year_of_capture + nb_eggs * nb_Chicks * Plsm_lineage * age * sex * SMI_Winter | year_of_capture + nb_Chicks + SMI_Winter | glm {n} |
| SMI in autumn | SMI | 13 | 36 | year_of_capture + nb_eggs * nb_Chicks * Plsm_lineage * age * sex | 1 | glm {n} |

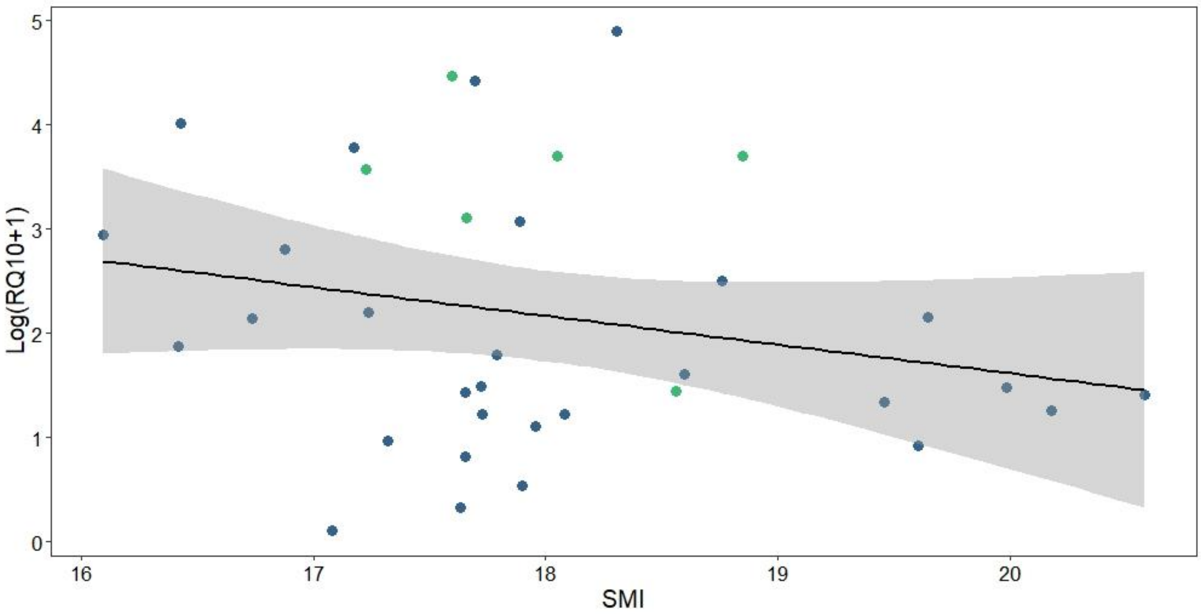


**Figure S1** – **Relationship between the scaled mass index (SMI) and the parasitemia (Log(RQ10+1)) of birds captured in autumn.** Blue dots correspond to individuals infected with *P. relictum* SGS1, green dots correspond to individuals infected with *P. homonucleophilum* SW2.
